# A thermo-responsive plasmid for biconditional protein expression

**DOI:** 10.1101/289264

**Authors:** Agathe Lermant, Alicia Magnanon, Alexandra Silvain, Paul Lubrano, Marie Lhuissier, Camille Dury, Maryne Follenfant, Zoe Guiot, Gaëtan Christien, Pauline Coudert, Antoine Arvor, Maxime Sportich, Gabrielle Vuillaume, Nicolas Delettre, Julie Henry, Eliott Lafon, Thomas Lhernould, Fanny Richard, Alexandre Ismail

## Abstract

Here we develop a temperature sensitive expression vector that allows the selective production of distinct proteins over different temperature ranges with a single plasmid. We use the *E. coli* cold shock translational control system (the cspA 5’Untranslated Transcribed Region - UTR) to drive the expression of a desired protein below a certain temperature threshold, and the lambda phage pL/cI857 transcriptional repressor system to drive the expression of a different protein above a certain temperature threshold. In this developmental work we use the chromogenic reporter proteins amilCP (blue chromoprotein) and monomeric red fluorescent protein (mRFP) to assess the function of the thermo-sensitive regulatory sequences over the desired temperature ranges. Our results show temperature dependent response of the cold shock regulatory sequence. However, our sequence design for the heat shock regulatory sequence did not give the intended result. The integration of these two temperature sensitive elements into a single plasmid awaits the re-design of the heat shock sequence.

## Introduction

### Motivation

Climate change is increasing the severity and frequency of high and low temperature extremes. [REF 0]. Departures from normal temperature ranges in agricultural regions has been responsible for major agricultural yield losses around the world [REF 1].

Therefore, the development of a low-cost thermal crop protection system is of great value to maintaining agricultural output in the current context of climate change. Here we describe the development of a thermo-responsive expression vector that allows the selective production of different protective compounds over different temperature ranges with a single plasmid. This work also demonstrates the value of molecular modeling in the rational design of genetic control sequences. Molecular modeling was used extensively in the development of the cold shock component and is likely to benefit the improvement of the heat shock component.

Departures from normal temperature ranges in agricultural regions can subject crops to conditions outside of their growth or even survival ranges. Outside of the semi-controlled conditions of agriculture, many organisms live in environments where temperatures can exceed the freezing and boiling points of water, which is essential to life. Many of these organisms, such as micro-organisms, insects, and plants, have little or no ability to move in response to temperature extremes. They have consequently evolved molecular mechanisms for sensing temperature extremes and coupled them to genetic responses to adapt cellular processes to these physical challenges. These are known as the cold shock and heat shock responses, whose biological bases have been well described in the literature. [REF 5**] [**REF 18] In this project we use temperature sensitive genetic sequences from known cold shock and heat shock systems to induce protein expression only above or only below certain temperatures thresholds.

### Cold responsive system

The survival and growth of *E. coli* can be significantly impaired when subjected to cold (near freezing) shocks or sustained cold temperatures. Membranes lose fluidity, global enzyme activity decreases, and mRNAs tend to form secondary structures which impair ribosomal translation [REF 3]. To counter the effects of cold temperatures, cells express cold shock proteins (CSP).

CSPs help translation to proceed at low temperatures [REF 4]. They are a large family of nucleic acid chaperone proteins whose expression is significantly up-regulated in response to cold stress [REF 2]. The *E. coli* Cold Shock Protein A (CspA) and its upstream regulatory sequences is one of the most well studied examples [REF 5] and forms the basis of our cold-responsive expression control sequence.

Cold induced protein expression by the CspA promoter has both a transcriptional and translational basis, although some details are still unclear. CspA transcription has been shown to increase up to three-fold relative to its level at 37°C [REF 6]. CspA promoter activity is additionally enhanced by an upstream sequence motif called the UP element [REF 7**] [**REF 8]. CspA expression is also controlled at the translational level through at least two molecular mechanisms. At temperatures near 37°C CspA messenger RNA (mRNA) is actively degraded by an RNAse and has a half-life of 12 seconds, making it essentially unavailable for ribosomal translation [REF 6]. At temperatures near 15°C and lower the RNAse is inactive and CspA accounts for nearly 10% of cellular protein production [REF 10**] [**REF 11**] [**REF 12]. The CspA mRNA is likely stabilized through the formation of specific secondary structure motifs that are stable only at low temperatures. In this work we focus on the translational control system to apply it to other proteins of interest.

The translational control system comes from the CspA 5’UTR, an unusually long and conserved sequence common to all cold shock proteins [REF 3]. This region contains several sequence motifs important for low temperature protein expression, including the Cold Box [REF 7], the “UP element” (upstream sequence of the CspA promoter), and the DownStream Box (DS Box). The DS box is an unusual sequence motif in that it extends past the start codon and into the coding region of the expressed protein [REF 7], requiring careful placement of the DS Box sequence in the reporter or effector gene of interest.

Molecular modeling was used to rationally design the placement of the DS Box in the reading frame of our low temperature chromogenic reporter protein amilCP. Further molecular modeling in this work strongly suggests that the DS Box contributes to the formation of a specific 5’UTR mRNA secondary structure which is recognized by <u>ribosomal protein S1</u> in the formation of the ribosomal pre-initiation complex [REF 13**] [**REF 14**] [**REF 15].

The UP element, CspA promoter, 5’UTR, and DS Box work synergistically to promote conditional low-temperature expression [Figure 1: Wild type CspA genetic structure]. Therefore, our design retains all of these regulatory sequence features as a control system for expressing the gene of interest. [Figure 2: Proposed genetic construction for cold-specific gene expression].

**Figure 1:**
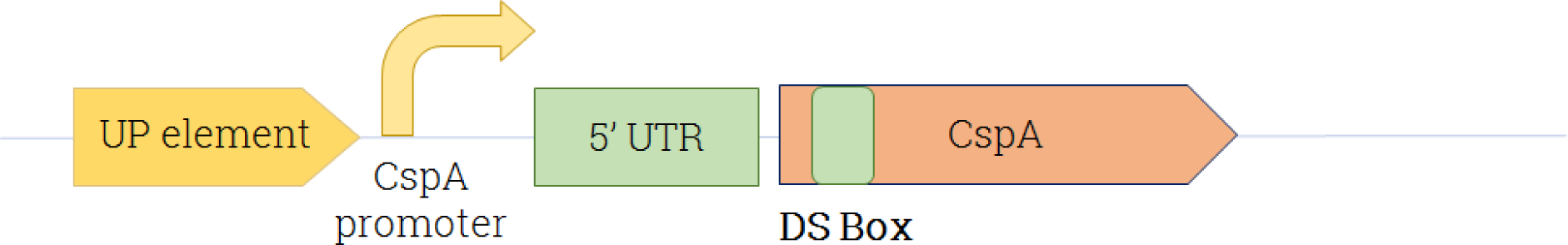
Wild type CspA genetic structure

**Figure 2:**
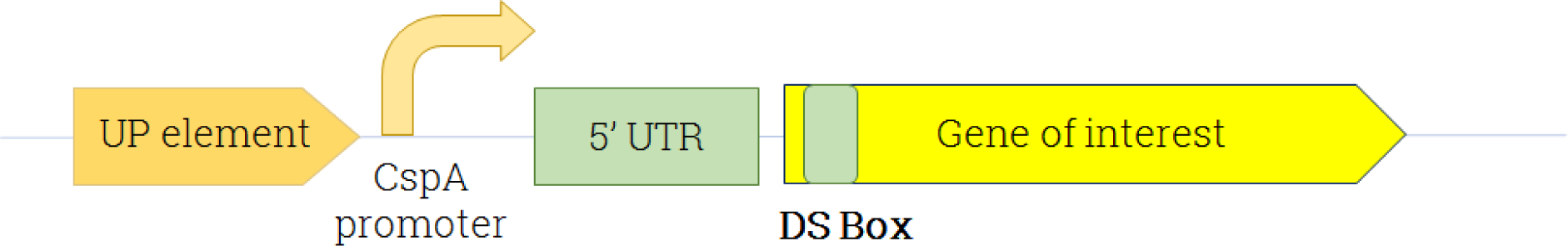
Proposed genetic construction for cold-specific gene expression

### Heat responsive system

The lambda phage of *E.coli* contains a well-studied transcriptional control system for heat induced conditional protein expression: the pL (or pR promoter) and its λcI repressor [REF 9]. Originally, these components serve as the genetic switch between the lysogenic and lytic phases of the phage life cycle. The creation of the cI857 repressor mutant of λcI made it thermo-labile above 37°C [REF 17], meaning repression becomes inactive above a certain temperature. This system is widely used for industrial production of recombinant proteins in *E. coli* under up-shift in temperature [REF 16].

Our strategy to induce the expression of any gene of interest at high temperature only will therefore be to create a recombinant plasmid containing the sequence coding for our reporter gene inserted downstream the pL promoter in presence of the cI857 repressor (Figure 4).

**Figure 3:**
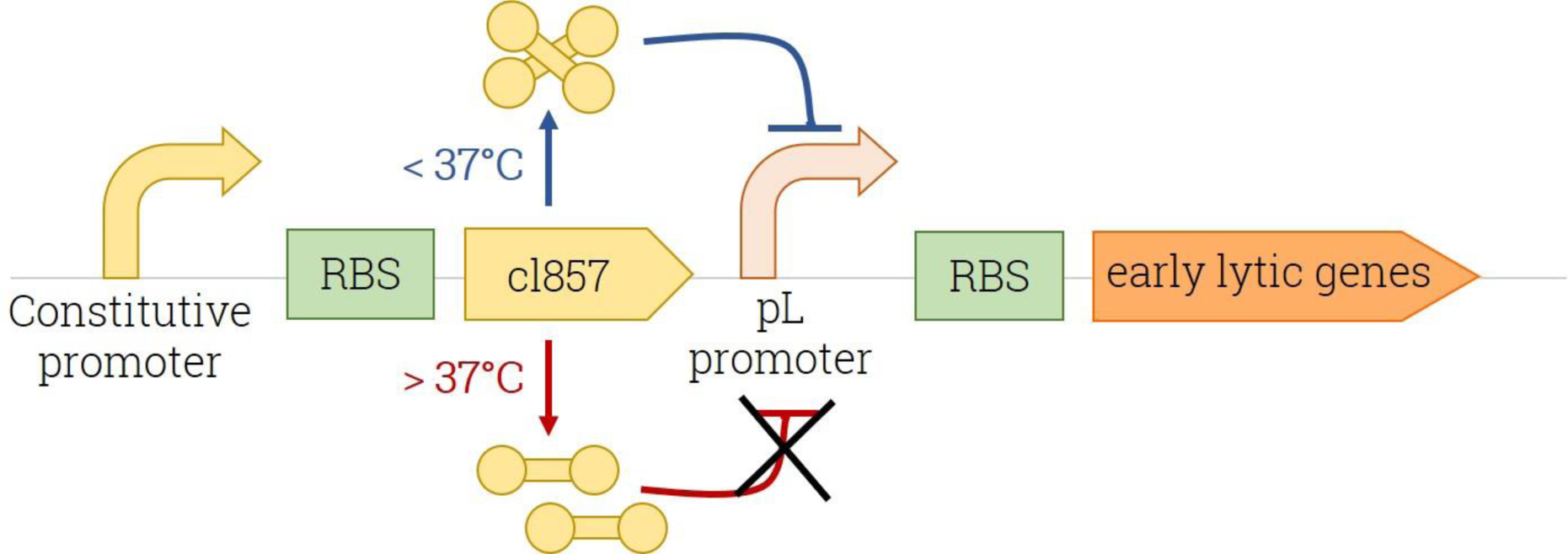
Representation of the pL/pR promoters controlled by the cI857 repressor [REF 18]

**Figure 4:**
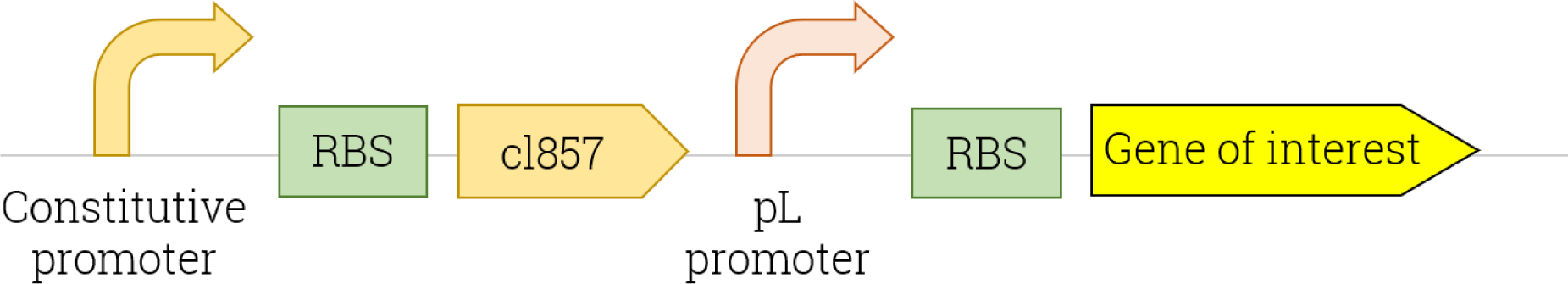
Proposed genetic construction for heat-specific gene expression

Ultimately, we intend to combine both the cold and heat responsive systems into a single construct for a biconditional thermo-responsive plasmid for expression of plant-protecting compounds upon departure from temperatures compatible with a given crop.

## Materials and Methods

### Chassis

*E. coli* was used as a chassis for plasmid construction because its genome is well studied and has several strains useful for molecular and cellular biology studies. The DH5α and BL21 strains were supplied by NEB (New England Biolabs) through the partnership with the iGEM Foundation. The DH5α strain was used for plasmid construction and cloning procedures because of its high transformation efficiency. The BL21 strain was used for expression assays because of its high level of protein expression. All our constructs were codon optimized for *E.coli* DH5α and verified through sequencing.

### BioBrick naming

The genes and genetic constructions used for the iGEM are commonly called “BioBricks”. A BioBrick is any sequence of DNA (promoter, Ribosome binding site, protein sequence, terminator…) that is flanked by a prefix (EcoRI and XbaI restriction sites) and a suffix (SpeI and PstI restriction sites). Every BioBrick shares the same prefix and suffix, allowing for efficient assembly.

The common naming of BioBricks in the iGEM registry is “BBa_x” where x is any identifier for the part in the registry. This naming strategy was used in this paper to refer directly to the part submitted to the registry by our team.

The registry is accessible here: http://parts.igem.org/Main_Page

### Plasmid backbones

Designed sequences were constructed with the <u>pSB1C3</u> and <u>pSB1A3</u> plasmid backbones. Final constructions were delivered on the pSB1C3 per iGEM Parts Registry standards. Plasmid pSB1A3 was used for the propagation of BBa_K2282011 (cold-response construct) which permits conditional low temperature expression of amilCP. The pSB1A3 backbone includes the ampicillin resistance gene and pSB1C3 includes chloramphenicol resistance and a high copy replication origin, which allows a high copy number per cell, facilitating DNA purification.

### Primer design

Primers were designed using the BioBrick suffix and prefix from the pSB1C3 backbone. Using Geneious 10.07 [REF 19], we extracted the sequences containing the EcoRI and PstI restriction sites in a reverse form from the BioBricks. The sequences had a size between 20 and 30bp with a Melting temperature (Tm) around 50°C and 60°C and a Guanine/Cytosine (GC) ratio of 50%. The sequences were then converted into primers. A “TATATA” sequence was added at the beginning of the forward primer and at the end of the reverse primer for DNA polymerase binding. All of our sequences were designed using Geneious by adding single BioBricks of DNA to form the intended DNA construct. Each sequence is formed by a BioBrick prefix and suffix flanking the sequences to which the primer binds. Each element was then introduced (Promoter - Ribosome Binding Site (RBS) - Gene of interest-terminator). Geneious was used to detect the restriction sites, design our primers, choose the best Tm for polymerase chain reaction (PCR), and visualize our system while adhering to the pSB1C3 backbone assembly standard.

### Molecular modeling software

Molecular modeling was used to rationally design the placement of the downstream box (DS box) inside the amilCP coding sequence for the cold shock system sequences (BBa_K2282011) and (BBa_K2282006). Molecular modeling was also used to explore the structural basis of cspA 5’ Untranslated transcribed region (UTR) function. Phyre2 [REF 20] was used to generate a 3D model of wild type amilCP and the DS box designed mutation. Phyre2 was also used to generate a 3D model of domain 3 of wild type ribosomal protein S1. SimRNA [REF 21] was used to generate 3D models of a 187 base-pair stretch of the cspA 5’UTR mRNA, including the downstream box present in the coding region. GROMACS [REF 22] was used to generate all atom XTC format trajectory files of SimRNA trajectories for visualization and analysis in VMD [REF 23]. ConSurf [REF 24] was used to identify the likely mRNA binding site of ribosomal protein S1 domain 3. Autodock Vina [REF 25] was used to perform a global blind docking of small model peptides from the DS box sequence onto the 3D model of ribosomal protein S1 domain 3. PatchDock [REF 26] was used to perform rigid body docking of the mRNA structure (from SimRNA) with the homology model of ribosomal protein S1-domain 3 (from Phyre2). PyMol [REF 27] and DiscoveryStudio [REF 28] were used to visualize selected results and prepare images.

### Design of sequences

Sequences were ordered and constructed by Integrated DNA Technologies (IDT) from the partnership with the iGEM Foundation of Boston. We imported the sequences from Geneious to IDT. IDT has to test the complexity of the sequences before synthesizing it. The sequence BBa_K2282013 had to be modified from its first draft due to its repetitive sequence content (arising from two double terminators). We integrated the last part of the sequence in a reverse form and we added a different terminator from the first part of the sequence to avoid concatemers. All the designed parts and their current status regarding the iGEM Parts Registry are summarized in Table 1.

**Table 1.** ID and description of the used designed parts

| Part reference | Part components |
| --- | --- |
| BBa_K2282005 | Anderson constitutive promoter (BBa_J23100) – amilCP |
| BBa_K2282006 | Anderson constitutive promoter (BBa_J23100) - amilCP+DSbox |
| BBa_K2282011 | UP element - CspA promoter - 5'UTR - amilCP+DSbox |
| BBa_K2282012 | pL promoter - mRFP (BBa_E1010) |
| BBa_K2282013 | Anderson constitutive promoter (BBa_J23100) - cL857 repressor (BBa_K098997)<br>- pL promoter - mRFP (BBa_E1010) |

### Cloning strategy

All sequences were digested with EcoRI-HF and PstI from NEB with a final concentration of 40U/µL and the digestion mix was incubated for two hours at 37°C. We used the same protocol for the pSB1C3 backbone. Each digested sequence was ligated with digested pSB1C3 using T4 DNA ligase from NEB with a final concentration of 40U/µL. Competent *E.coli* DH5α were transformed with the ligation product into Super Optimal Broth (SOC) medium, using the classic heat-shock technique, and then spread on solid Lysogeny Broth (LB) medium with chloramphenicol and incubated at 37°C for 24 hours. Constructs were checked by colony PCR. Colony PCR is a protocol permitting the extraction of the total DNA of any transformed bacteria and the amplification of its plasmid DNA with primers to visualize if the bacteria have included the right plasmid DNA.

### Part characterization: colorimetric measurements over time

#### Visual color test

Cold and heat response were tested through colorimetry. The test was run over three days. Because their growth rate is slower in cold temperatures, bacteria were first pre-incubated at 37°C until reaching an OD600 of 0.5, corresponding to the exponential phase. Cultures were then split in half and held at temperatures according to the genetic construction to be tested:

- For the Cold-Response system, one half at 12°C/15°C/20°C/27°C and the other at 37°C
- For the Heat-response system, one half at 27°C and the other at 37°C

For each condition, a sample of 1 mL was collected after 20h and centrifuged at 13.000 g. This duration was sufficient to obtain similar OD600 values around 0.7. Incubation at 37°C was necessary to obtain a positive control. Other controls were used as well for the heat and cold-response system based on the constitutive expression of amilCP and mRFP (see BBa_K2282005 and BBa_K2282012). Incubation temperatures were chosen based on bibliographic studies [REF 3].

#### Absorbance measurements over time

After being transformed, bacteria were grown at 37°C for 60 hours and Optical Density (OD) at 588 nm was measured every 10 min approximately with a Spark 10M Tecan. OD 800 was taken as a bacterial growth indicator, as the commonly used OD600 was too close to our wavelength of interest and could therefore interact in the results.

#### Blue profile plots

Blue intensity profile plots of bacteria pellets were obtained using the ImageJ software. Rectangular areas were defined within each pellet obtained on the picture, the RGB intensity values were extracted from each pixel of the area selected and integrated into a profile plot. Red and Green intensities were discarded.

## Results

### Modeling results

#### amilCP homology modeling

The cold-sensitive regulatory sequence of this construct is the *E. coli* cspA 5’ untranslated region (5’UTR). The majority of this sequence is upstream of the start codon. However, the Downstream (DS) box motif is present after the start codon and is translated as part of the target protein. Therefore, we studied the N-terminal region of amilCP (a blue chromogenic reporter protein) with molecular modeling in order to determine a DS box sequence placement that would preserve the structure and function of the protein.

While the structure of amilCP has not been experimentally solved, several other members of the family (coral chromoproteins which are GFP-like) have been crystallized and their coordinates deposited in the Protein Data Bank (PDB). Therefore, we decided to use homology modeling to create model of amilCP to understand the structure of the N-terminal region and the likely impact of modifications to this region.

The Phyre2 homology modeling server [REF 2] was used to find templates, generate alignments and create 3D models. Phyre2 was selected because of its hidden Markov Model-based fold library, which allows detection of remote sequence homologs down to ~15% sequence identity. In the case of amilCP, this search revealed a homologous coral chromoprotein (PDB entry 1MOU) with 93.2% sequence identity (as shown in the template-target alignment in Figure 5), permitting the construction of a reliable model.

**Figure 5:**
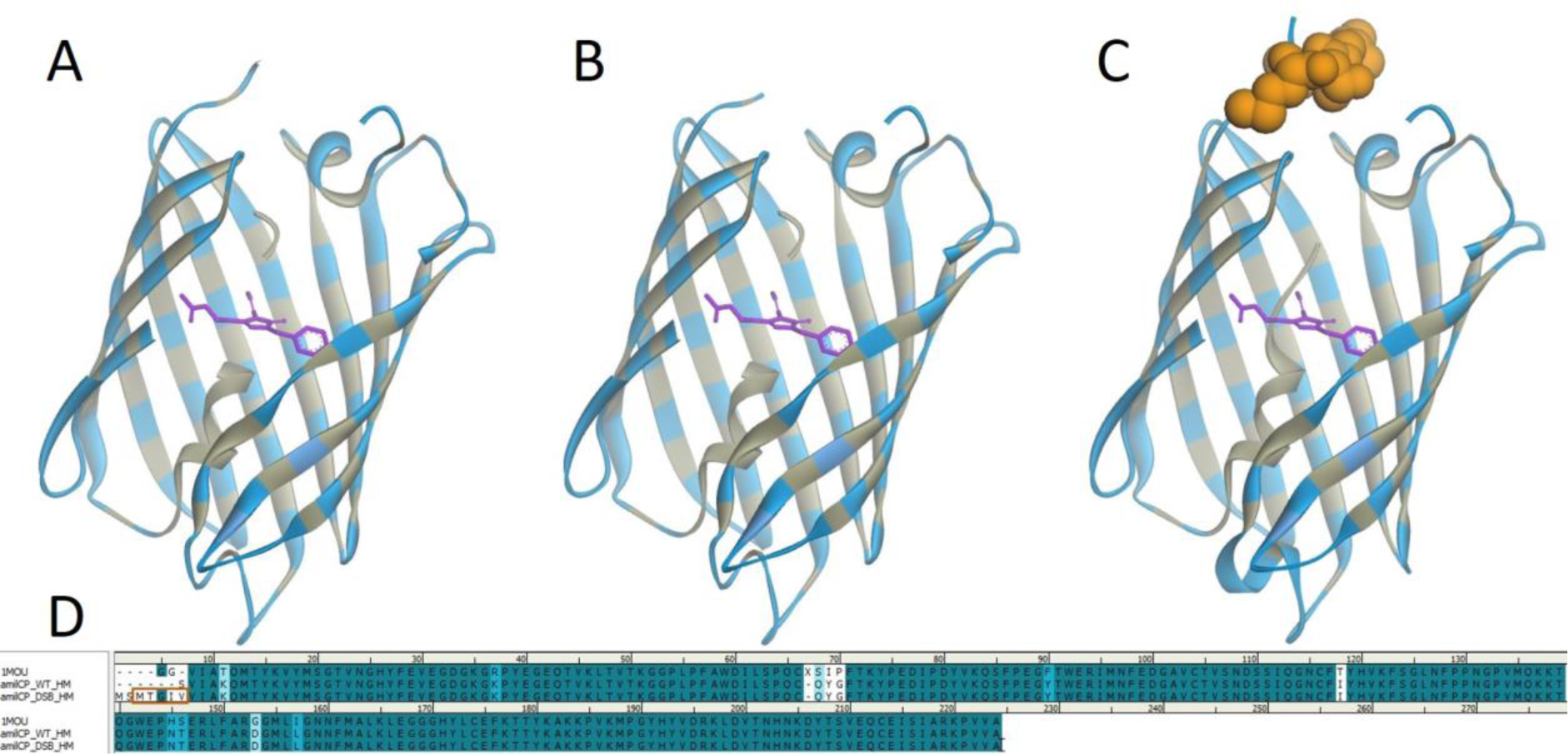
Sequence alignment and 3D structures of 1MOU, Homology Model_WT, Homology Model_DSB

**Figure 6:**
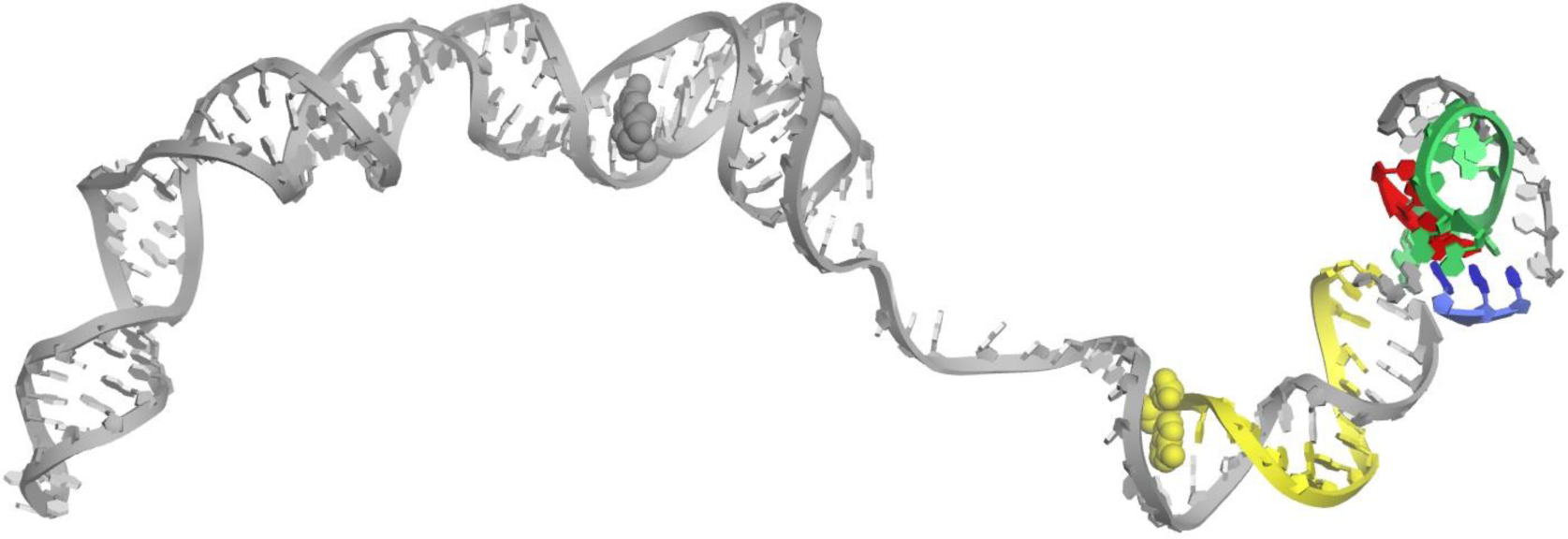
iGEM 187 mRNA structure computed by SimRNA. The SD sequence is in red; the DS box is in yellow; the putative S1 protein binding sequence is in green; the start AUG codon is in blue.

Figure 5 shows the highly similar structure between the template (PDB structure 1MOU) and the Phyre2 homology model with an RMSD of 0.232 angstroms over identical residues. The first six N-terminal residues (GGVIAT of 1MOU) does not display backbone angle values describing a regular secondary structure element and is predicted to be solvent accessible, implying that it does not form essential structures or contacts with the rest of the amilCP structure.

Based on these structures, we decided to insert the DS box sequence MTGIV (AUGACUGGUAUCGUA) after the first two residues of the wild type amilCP sequence.

We then made a homology model of this mutant amilCP sequence with Phyre2 in order to assess the likely position and structure of the additional residues relative to the amilCP beta barrel, as shown in orange in Figure 5 above.

Based on these results, we decided to proceed with the synthesis of this mutant sequence IDT and plasmid construction.

#### cspA 5’UTR structure modeling

The cspA 5’UTR is known to exhibit the following behaviors which drive conditional expression as a function of temperature. At low temperatures it enhances transcription by binding to ribosomal subunit 1 through the S1 protein [REF1], whereas at high temperatures it negatively regulates expression by favoring RNAse mediated degradation of its transcript [REF2].

The cspA 5’UTR has been partially structurally characterized by the use of the Tn5-lac transposon insertion [REF3] and gel retardation analysis [REF4]. The general hypothesis is that the cspA 5’UTR mRNA assumes a specific secondary structure at low temperatures. However, experimental characterization of this conformation thus far has not yet proposed a 3D structure for this mRNA sequence, leaving ambiguity concerning structure-function relationships of various other CspA 5’UTR sequence motifs such as the Cold box and DS box.

In order to get a better idea of possible structure-function relationships for rational design we used molecular modeling to create structural models of the 3’ end of our cold shock construct mRNA (a 187-nucleotide segment). This includes the DS Box, a key sequence motif of the cspA 5’UTR mRNA.

SimRNA (an ab initio coarse grained replica exchange Monte Carlo method) was used to generate an ensemble of conformations covering a folding trajectory of the given mRNA sequence. The ensembles converge to a set of similar structures, revealed through all vs all root-mean-square deviation (RMSD) based clustering. The resulting top cluster centroid was selected as a representative conformation and retained for further study.

This portion of the mRNA is predicted to form an extended double helix structure, separated by a single stranded segment. The DS box sequence itself often forms a double helix structure, as it is present in 27 out of the 39 cluster centroids. This suggests that the DS box sequence may participate in a conserved structural motif important for low temperature transcription initiation - and one that involves other known conserved sequence motifs.

Review of the literature suggests that assembly of functional ribosomes at low temperatures depends on binding of mRNA to the small subunit in the pre-initiation complex, mediated by ribosomal protein S1 [REF 5]. In order to explore this, we pursued rigid body docking with a homology model of this protein.

#### cspA 5’UTR - S1 protein interaction modeling

We found from the literature that the Subunit 1 (S1) of the ribosome was necessary for the binding of RNA constructs [REF1] and more specifically for the initiation of the RNA-ribosome complex. This applies to many mRNAs, especially for those with none or weak Shine-Dalgarno sequences, and those with highly structured 5’ regions. More specifically, the domain 3 of S1 (S1-D3) was very likely to bind first to the 5’UTR of our protein(RNA) construct[REF2].

In order to explore the hypothesis of an interaction between the 3’ region of the temperature sensitive 5’UTR and ribosomal protein S1 domain 3 we decided to pursue docking of structural models of each.

Since there is no crystal structure of S1 domain 3 we searched for a template using the Phyre2[REF3] server. The model sequence was re-aligned against the top template using MAFFT-L-INSI [REF4], followed by I-TASSER[REF5] to build a final 3D model. On this structure, we observed a specific motif with conserved residues (as calculated by ConSurf [REF]) that we will call “valley” from now onwards. This valley seemed to be an externally accessible site, being located on the exterior side of S1-d3, while having two strands of highly conserved residues bordering it. As such, we considered it a likely binding site for our cspA 5’UTR mRNA structure.

As a reference, we compared it to a sequence found in the Protein Data Bank (PDB). This sequence, referred as the code 4v9r [REF6] on the PDB, consisted of S1-d3 bound to single stranded RNA (ssRNA) (as seen in Figure 7 panel A). We applied the same steps used for our homology model of S1-d3. On this structure, we also observe the same valley motif with highly conserved residues, reinforcing our previous hypothesis.

**Figure 7:**
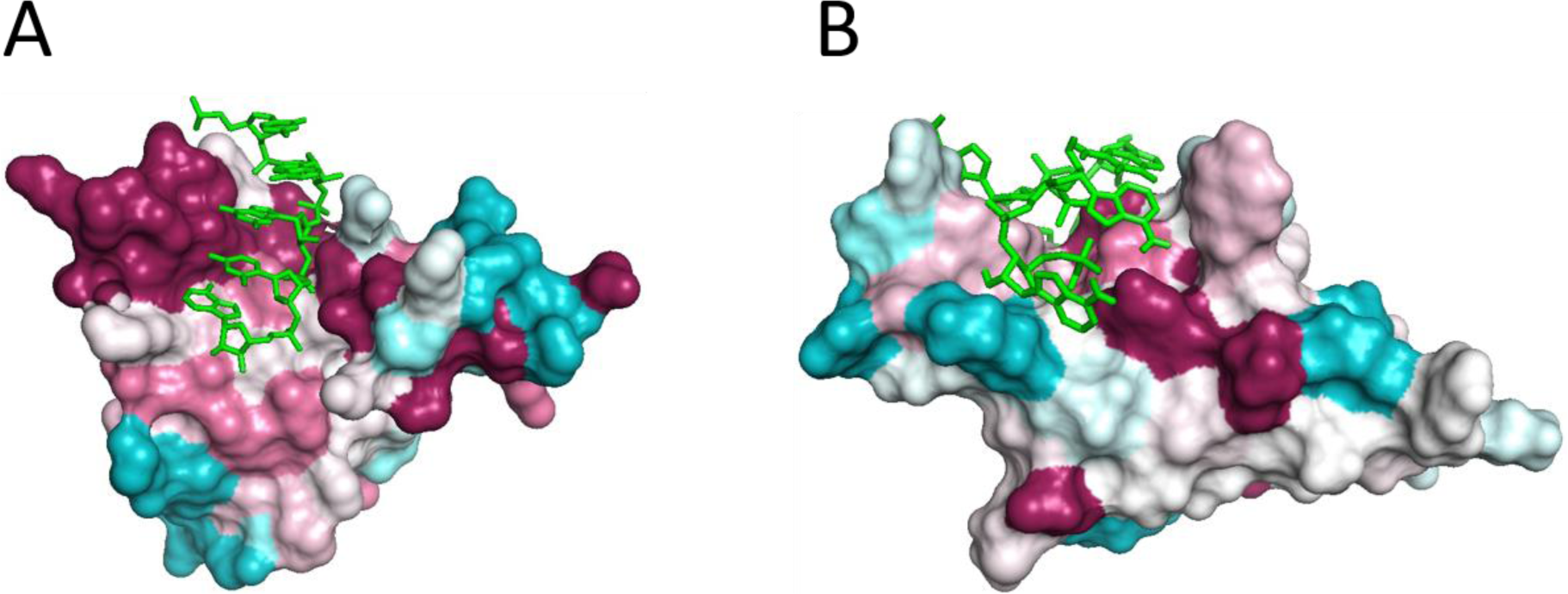
A) *Thermus thermophilus* S1-d3 (colored by residue conservation) complexed with ssRNA (green) from PDB entry 4v9r (X-ray structure). B) *E. coli* S1-d3 homology model colored according to residue conservation docked with DS Box ssRNA (green).

**Figure 8:**
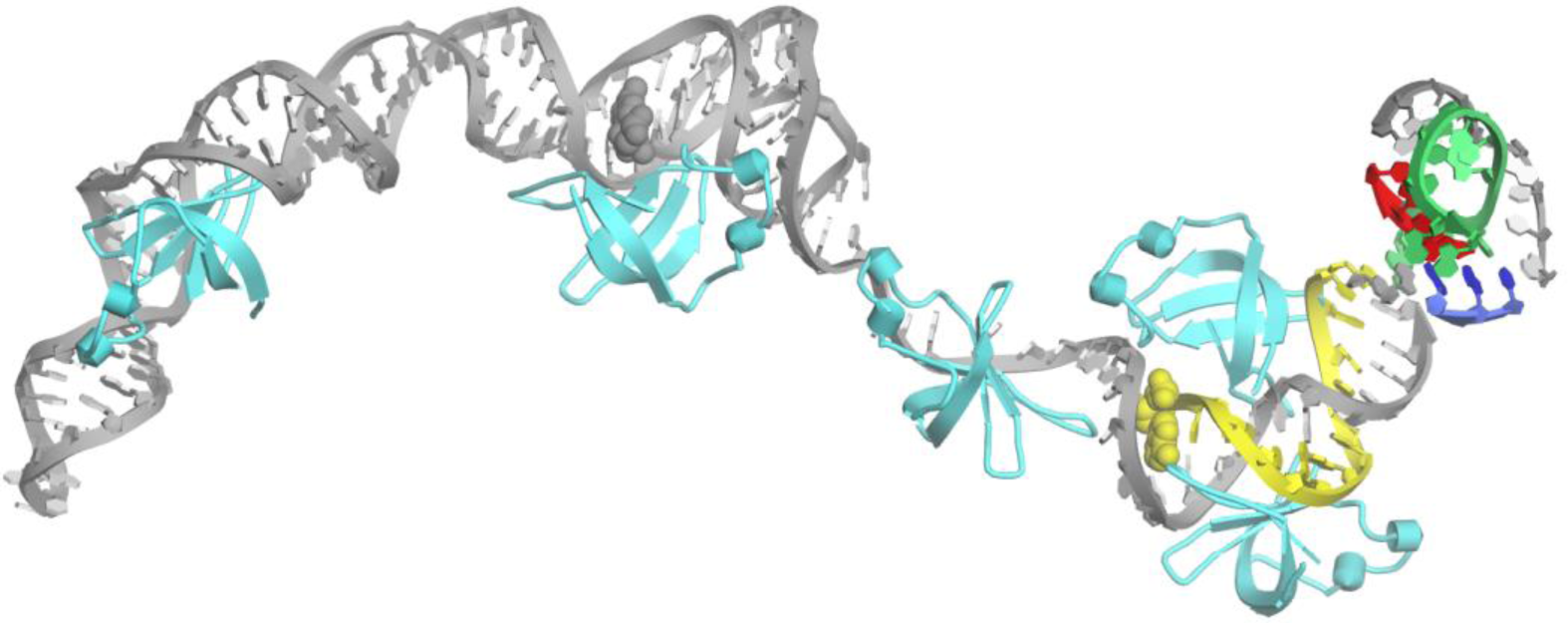
S1-d3 (cyan) and cspA 5’UTR mRNA (grey) docking results. The SD sequence is in red; the DS box is in yellow; the putative S1 protein binding sequence is in green; the start AUG codon is in blue.

**Figure 9:**
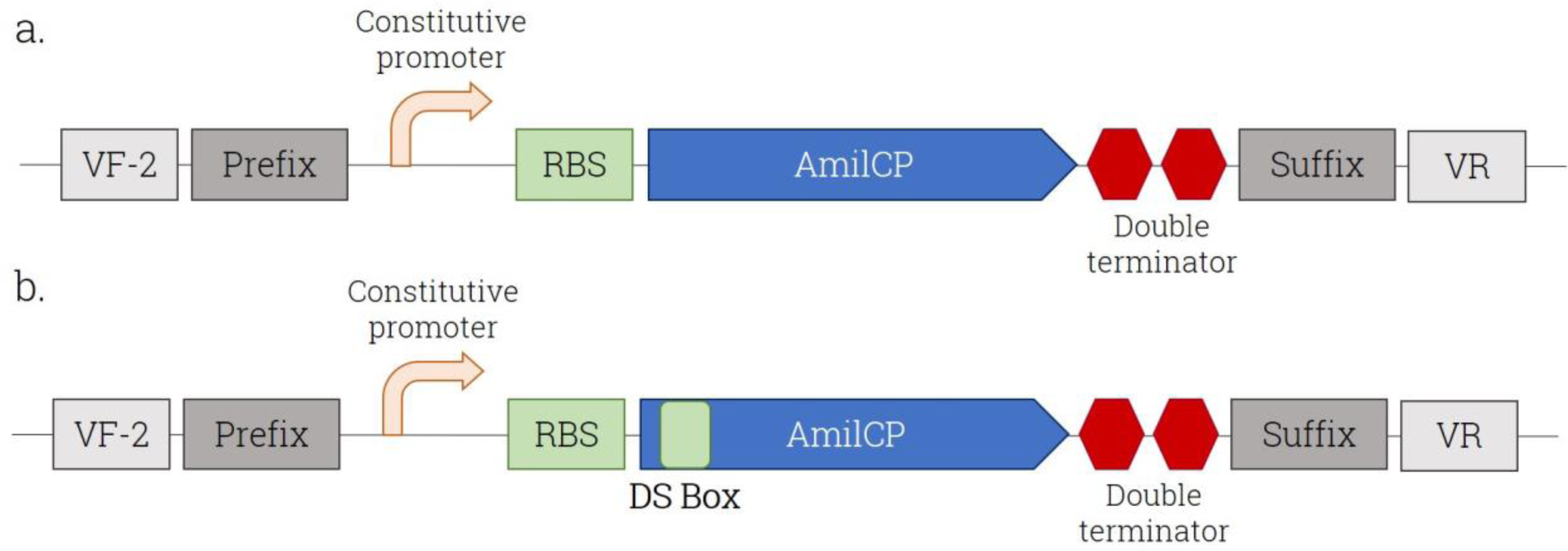
Genetic structures of the designed parts coding for the constitutive expression of Wild-type (WT) amilCP with/without the DS Box. **a.** Part BBa_K2282005 coding for WT amilCP; **b.** Part BBa_K2282006 coding for DS Box-amilCP

From the previous models, we decided to study more precisely the conservation of residues to determine the best docking site possible and test our valley hypothesis. This was done using the ConSurf web server [REF7] for both 4v9r and S1-d3. A globally blind docking by Autodock Vina was able to find poses at a site homologous to that seen in the 4v94 crystal structure.

These small-scale results were encouraging and so we proceeded to use our homology model of S1-d3 in docking with the larger mRNA structure previously computed with SimRNA. We used the PatchDock server[REF8], setting the receptor as S1_d3, the ligand as our 5’UTR (SimRNA cluster 1 result) and using default settings.

The top 20 results contain a set of interesting positions. First, we note that the DS box forms part of a double helix secondary structure which appears as a favorable binding site for our homology model of S1-d3. We also not that there are several other binding sites in approximately linear distribution on an extended mRNA double helix - indeed there appear to be room for 5 or 6 simultaneously bound S1 sub-domains, all of which have homology to domain 3.

## Experimental results

### Influence of the DS box on amilCP expression

#### 1. Impact of the DS box on amilCP color

We first needed to verify the expression and color properties of amilCP (blue chromoprotein) with the addition of the Downstream (DS) Box. Wild-type amilCP under constitutive expression was tested as Registry Part BBa_K2282005. DS Box-amilCP under constitutive expression was tested as Registry Part BBa_K2282006. The genetic structures of these constructs are presented in the figure below.

We observed the same color for both constructs, as confirmed through the absorbance spectra obtained (Figure 10). However, the DS Box version produced a slightly fainter color (Figure 11), which may be related to total amilCP expression levels.

**Figure 10:**
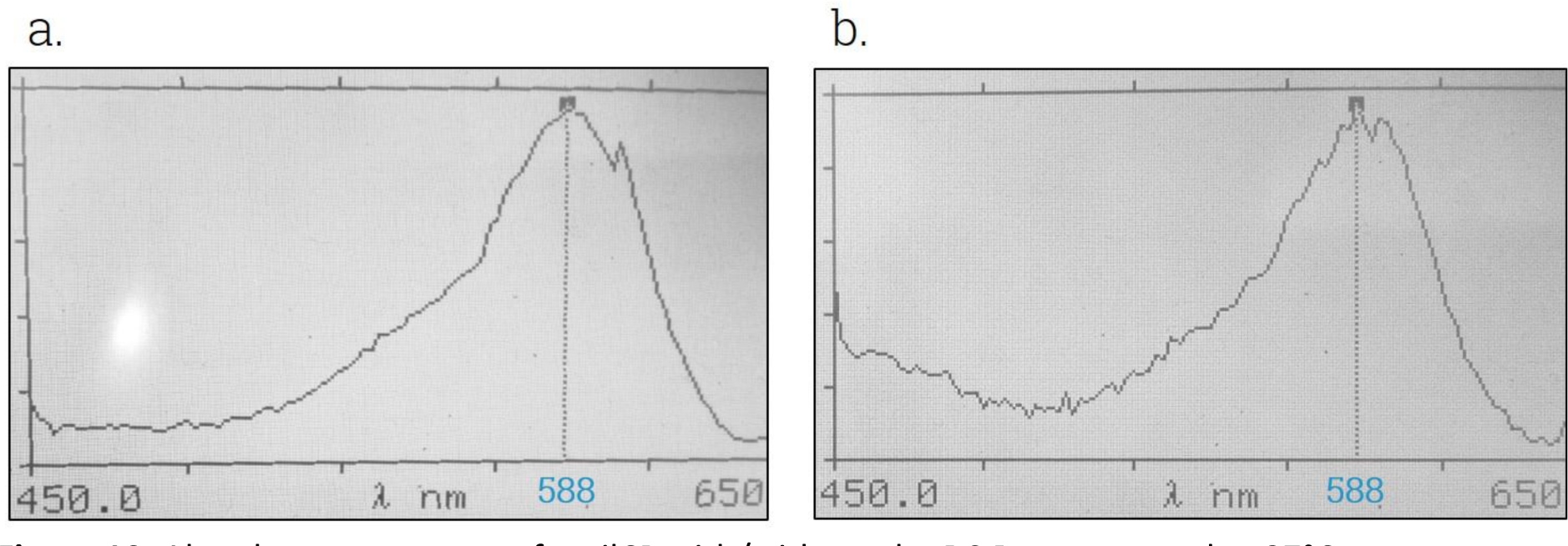
Absorbance spectrum of amilCP with/without the DS Box expressed at 37°C. **a.** WT amilCP from part BBa_K2282005; **b.** DS Box-amilCP from part BBa_K2282006

**Figure 11:**
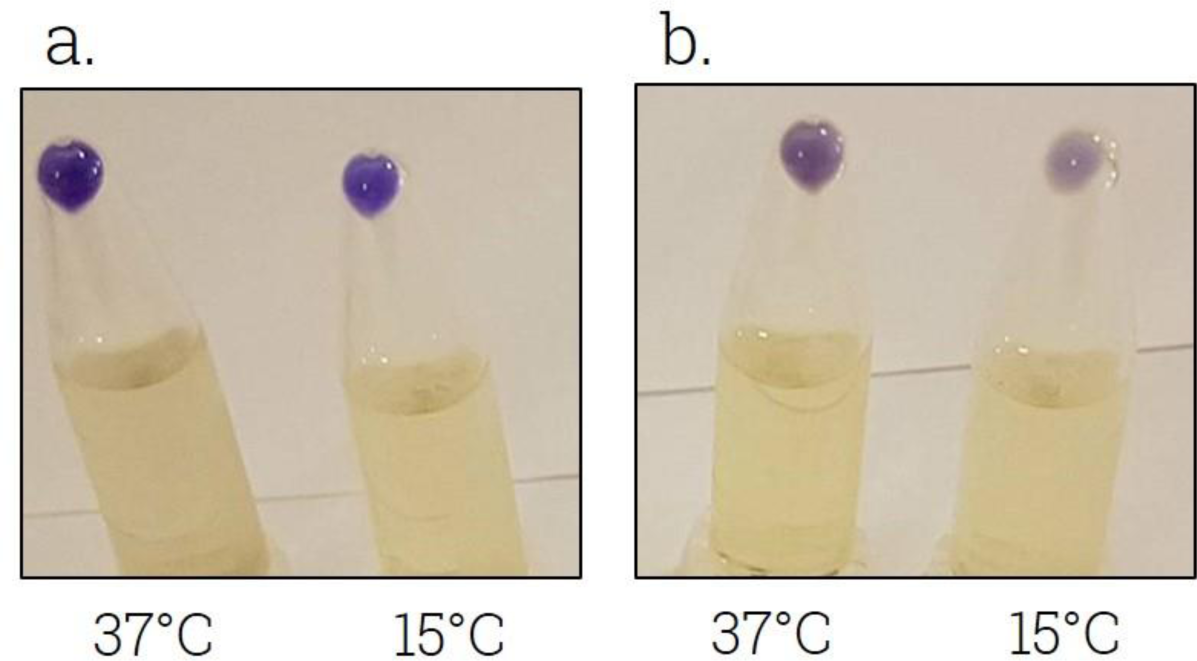
Color of amilCP liquid culture at 37°C and 15°C with/without the DS box *Pellets of E.coli DH5-α transformed with a. pSB1C3-BBa_K2282005 (WT amilCP) b. pSB1C3-BBa_K2282006 (DS Box-amilCP), and cultivated in liquid LB medium supplemented with chloramphenicol at 37°C and 15°C for 18 hours*

**Figure 12:**
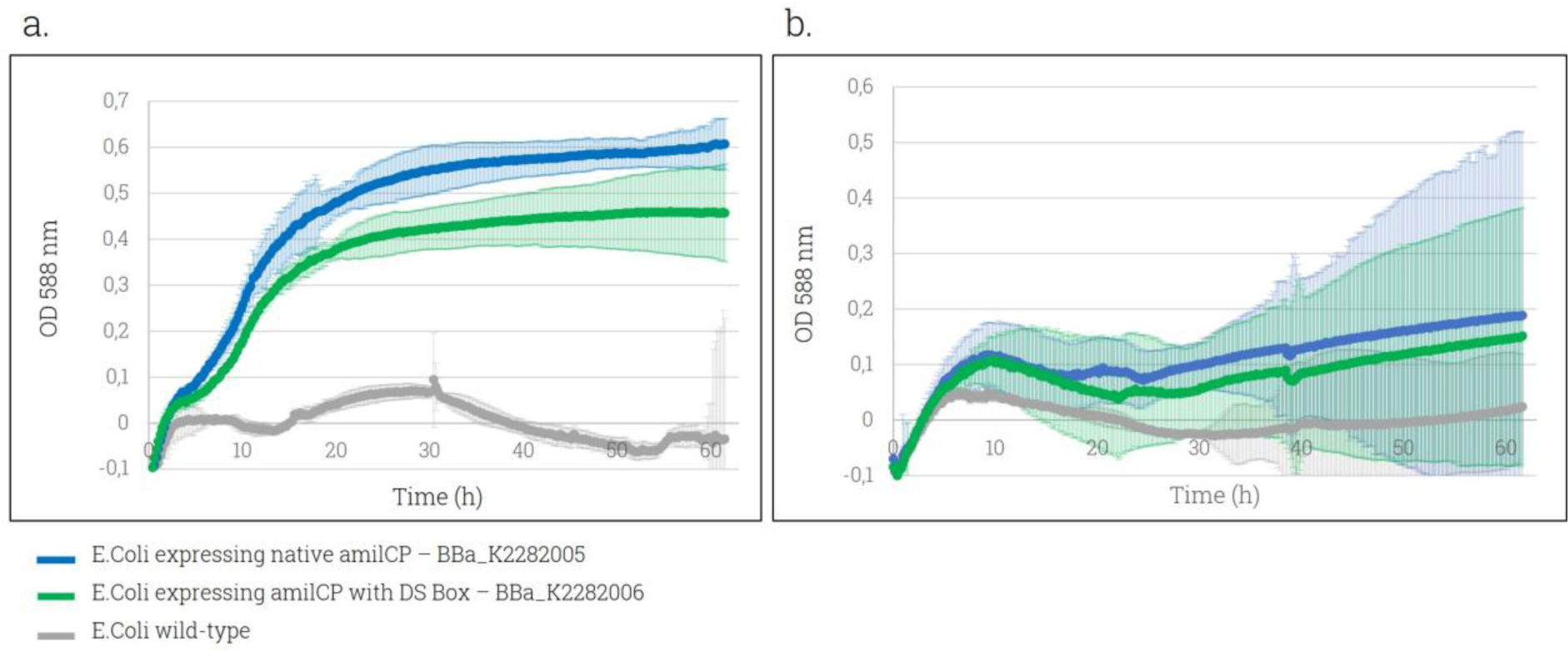
amilCP expression over time with/without the DS box at 37°C and 18°C **a.** OD588nm given by pSB1C3-BBa_K2282005 (WT amilCP) and pSB1C3-BBa_K2282006 (DS Box-amilCP) expression at 37°C; **b.** OD588nm given by pSB1C3-BBa_K2282005 (WT amilCP) and pSB1C3-BBa_K2282006 (DS Box-amilCP) expression at 18°C

#### 2. Impact of the DS box on amilCP expression over time

The impact of the DS box on amilCP expression level was evaluated at two different temperatures through the measurement of the OD588nm over time. The expression was first measured at 37°C, as it is the *E. coli* optimal growth temperature, then a lower temperature was chosen in order to assess the potential impact of the DS Box on the protein expression in cold conditions. 18°C was selected based on the material available.

The increasing of the signal at OD588nm over time for both transformed bacterial batches shows the expression of functional amilCP compared to the non-transformed ones.

This confirms that the presence of the DS box does not impact the protein absorbance properties.

The same protein expression pattern is observable at both temperatures, with an overall lower expression at 18°C. This result suggests the absence of enhancing properties on protein expression at 18°C from the DS box alone. Despite the presence of the protein in both cases, these curves also show a slightly better expression without the DS box, the difference being more pronounced at 37°C.

This result is in accordance with our hypothesis that the presence of the DS box influences the amilCP expression level, which may be related to the modification of amilCP mRNA folding.

Originally, the DS box is suggested to enhance mRNA translation during cold shocks [REF 29]. However, those results tend to show that without the whole cold-inducing machinery, the DS box does not seem to operate its enhancing action. Further characterization of our complete cold-shock plasmid showed its efficiency through a good protein expression at 15°C and not 37°C. This proves that a lower expression rate possibly induced by the presence of the DS box is not problematic when combined with the CspA promoter and 5’UTR sequence, allowing its activation.

### Test of the complete cold-responsive system

The complete cold-inducible construct was tested as Registry Part BBa_K2282011, whose genetic structure is presented in Figure 13.

**Figure 13:**
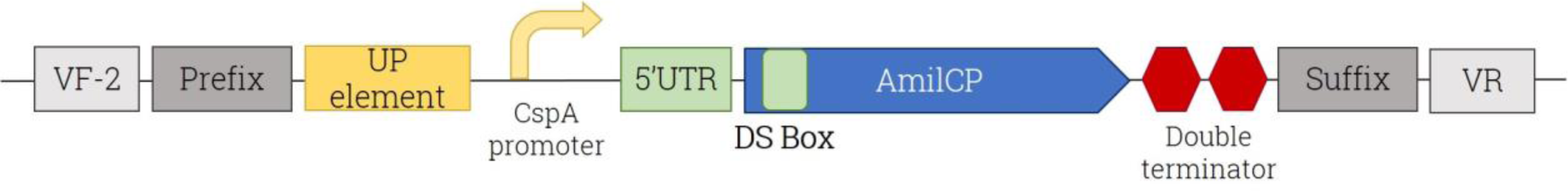
Genetic structure of the designed part coding for the conditional expression of amilCP at low temperatures (Part BBa_K2282011 in the iGEM part registry)

The construct was reported to induce a conditional expression under 15°C, and no expression above 37°C. Most of the previous experiments on cold-shock proteins were carried out only at 15°C and 37°C and the gene expression pattern between those limits is poorly known. In order to get a better insight of it, we evaluated the amilCP level over different temperatures ranging from 37°C to 12°C. Bacteria transformed with the cold-responsive construct were pre-incubated at 37°C until the OD at 600nm reached 0.5. They were then grown at different temperatures (12°C, 15°C, 20°C, 27°C, 37°C). Pellets containing an equivalent amount of bacteria were collected for each condition, based on similar OD600 measurements and equal culture volumes sampled. Their respective coloration was observed and compared as a function of the temperature gradient. Almost absent at 37°C, a blue color started to develop slightly at 27°C and appeared gradually higher as the temperature was lowered down to 12°C (Figure 14).

**Figure 14:**
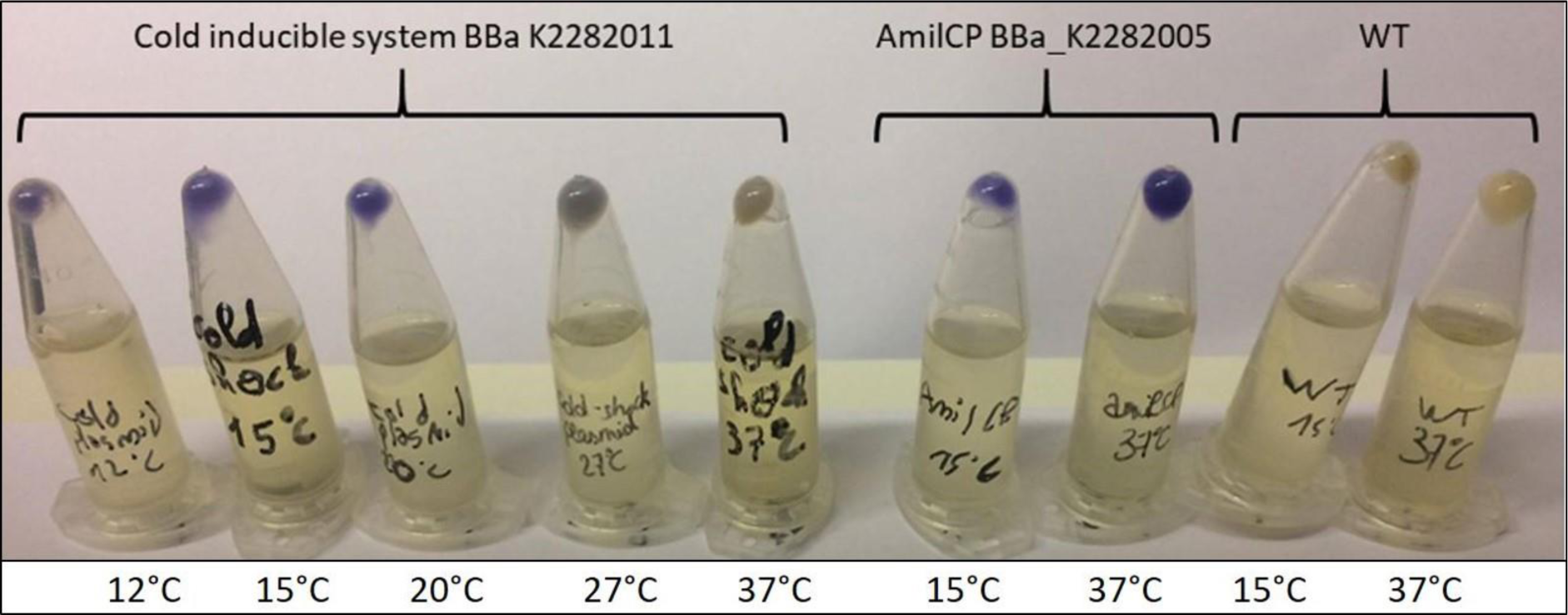
Visual aspect of amilCP expression gradient induced by the complete cold-responsive system from 37°C to 12°C. Pellets of DH5α transformed with BBaK2282011, after 20h of incubation at respectively 12°C, 15°C, 20°C, 27°C and 37°C (left), pellets of DH5α transformed with BBaK2282005 after 20h of incubation at respectively 15°C and 37°C (middle, positive control), and pellets of wild type DH5α (right, negative control).

Since equivalent bacteria amounts were collected here, we can consider that any difference in pellet coloration correlates with a differential amilCP expression per bacteria. Visual observations confirmed that our part BBa_K2282011 works as expected and allows efficient protein production at temperatures below approximately 20°C only, until low temperature as 12°C. Moreover, the cold response system blocks greatly but not completely the protein expression at higher temperatures above 27°C.

As we did not have a fluorimeter in our laboratory, amilCP was used as a reporter gene for observing the results with the naked eye. However, it was difficult to perform OD measurements without extracting proteins. Indeed, amilCP maximum of absorbance of 588 nm is close to 600 nm, the wavelength commonly used to measure the bacteria concentration. This proximity was prone to alter the actual signal corresponding to the amilCP protein. Measurements of OD588nm on extracted proteins would therefore constitute an interesting perspective to precise those results.

Despite this lack of equipment, we decided to complete these visual observations by a quantification of the blue value per pixel in each respective pellet obtained in Figure 14. The results are given in Figure 15:

**Figure 15:**
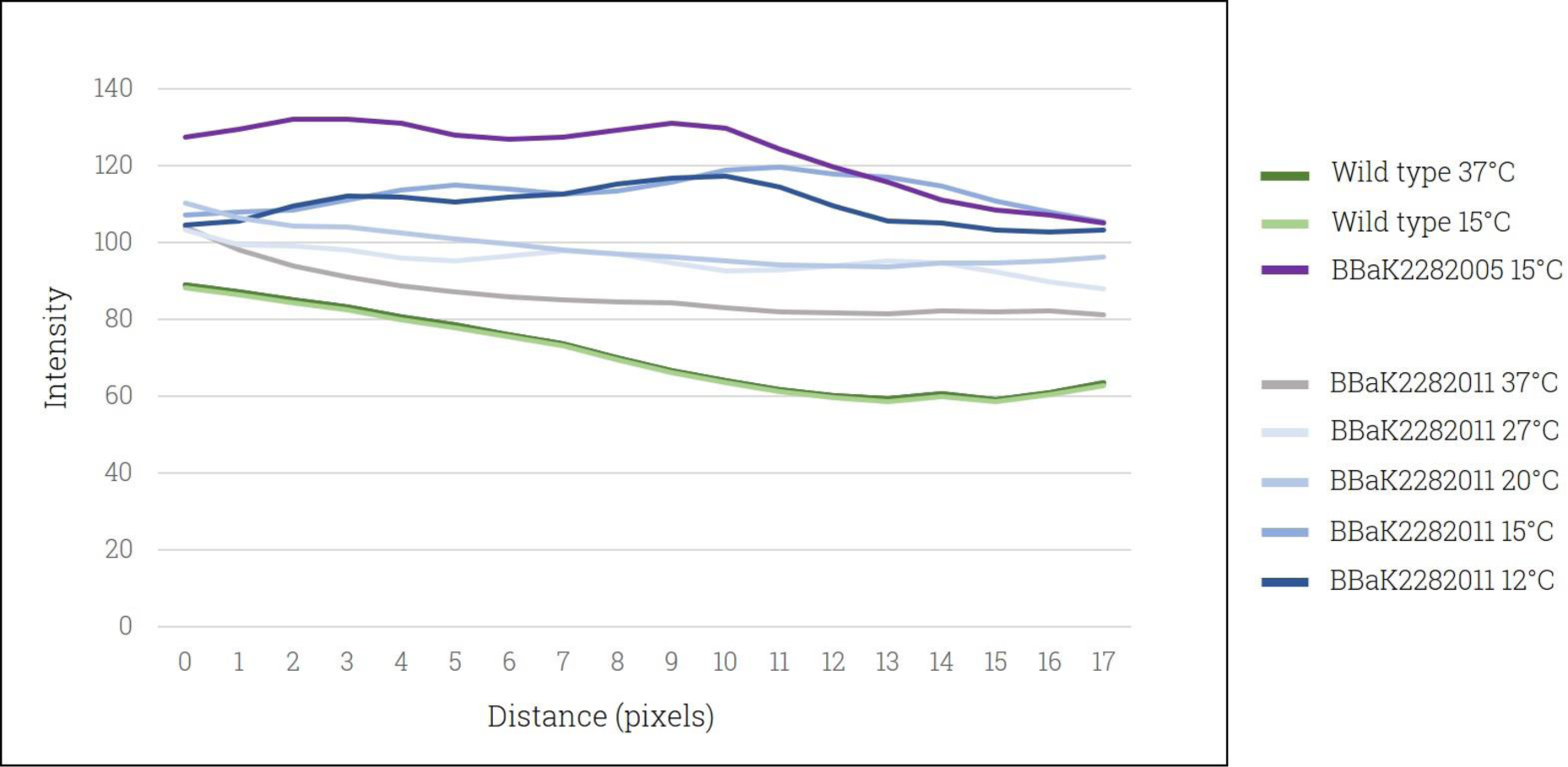
Blue profile plot of bacteria pellets expressing amilCP under the complete cold-responsive system from 37°C to 12°C.

**Figure 16:**
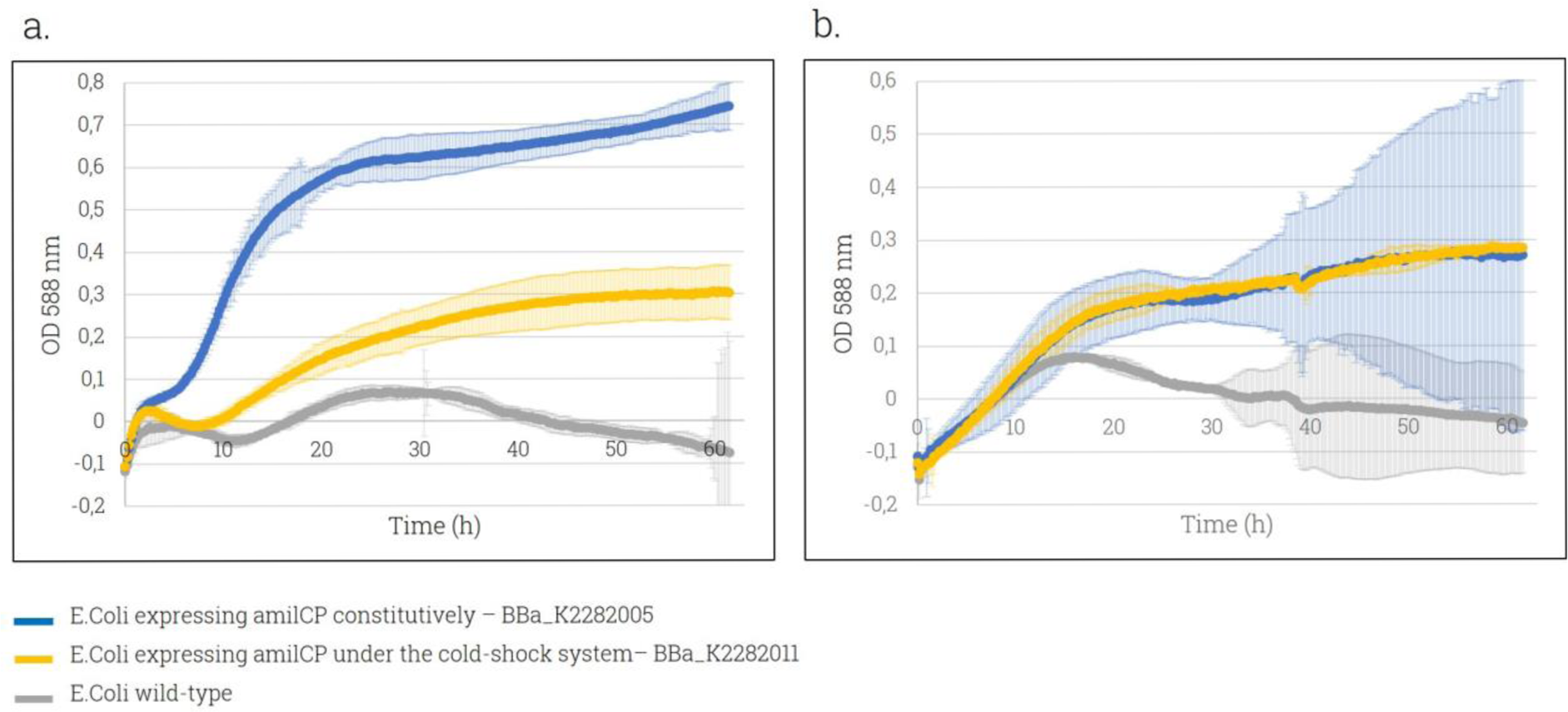
amilCP expression over time under the cold-responsive system at 37°C and 18°C **a.** OD588nm given by pSB1C3-BBa_K2282005 (constitutive amilCP expression) and pSB1C3-BBa_K2282011 (cold-induced amilCP expression) at 37°C; **b.** OD588nm given by pSB1C3-BBa_K2282005 (constitutive amilCP expression) and pSB1C3-BBa_K2282011 (cold-induced amilCP expression) at 18°C.

The blue intensity profile obtained for each pellet obtained was globally in accordance with our expectations, displaying a gradual increase as the temperature drops for bacteria transformed with the cold-responsive plasmid. At 37°C however, a slightly higher value compared to the negative controls suggest a small residual expression of amilCP at high temperatures and therefore the absence of a complete switch-off system. A significant gap was observed in amilCP expression between 20°C and 15°C, which appears to be the range where the cold-shock response mainly takes place. The intensity values obtained for the BBa_K2282005 positive control at 37°C appeared inconsistent with the intense coloration observed in the pellet, and where therefore not shown here. These results allowed us to precise our visual observations in absence of accurate equipment to properly extract and quantify proteins, however this method is prone to uncertainties when using a picture of medium quality, displaying pellets of unequal sizes.

According to our characterization results, the mRNA stability seems to increase progressively as the temperature decreases. This observation is in line with a degressive pattern more than an absolute expression switch-off system under a certain threshold. This is in accordance with the existing data on the CspA cold-shock system, which suggests a very short mRNA stability at high temperatures. It is then possible that a low translation still occurs during this time-lapse (1). Additionally, those multiple results were interesting in the sense that we did not find any previous data on amilCP expression between 15°C and 37°C.

### amilCP expression induced by BBa_K2282011 as a function of time at 37°C and 18°C

In order to follow the gene expression allowed by our cold-responsive genetic construct at high and low temperatures over time, *E.coli* BL21 were transformed with BBa_K2282011 coding for the cold-dependent amilCP expression. The amilCP expression was measured over 60 hours of growth at either 37°C and 18°C and the results were compared with E.coli transformed with BBa_K2282005 coding for constitutive amilCP expression (Figure 15).

At 37°C, the cold-responsive system induced a significantly lower amilCP expression compared to the control plasmid. This is in accordance with a conditional expression at low temperatures only. At 18°C however, the expression profile showed no difference between both constructions. Previous studies tended to show a significant expression under 15°C, however this temperature could not be tested due to the material available.

### Test of the complete heat-responsive system

The complete heat-inducible construct was tested as Registry Part BBa_K2282013, with the genetic structure is presented in Figure 17.

**Figure 17:**
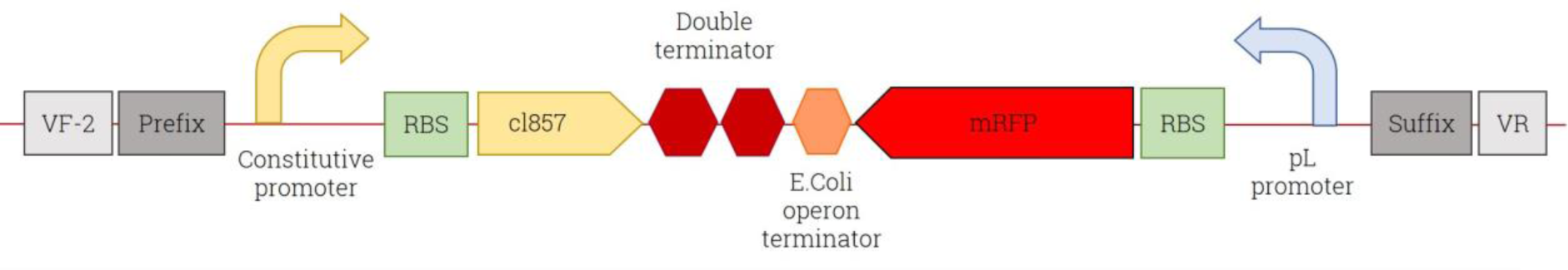
Genetic structure of the designed part coding for the conditional expression of mRFP at high temperatures (Part BBa_K2282013 in the iGEM part registry)

After bacterial transformation of DH5-α *E.coli* cells with the heat-shock and the positive control plasmids, we incubated them into Petri dishes added with solid medium at 30°C and 37°C. At 37°C, both bacterial populations appeared strongly red. Surprisingly, at 30°C, the populations also displayed a slight red color. Because of this temperature was not optimal for E.coli growth, the results were not conclusive. To get a better insight into the monomeric red fluorescent protein (mRFP) expression level at low and high temperature, we incubated our transformed bacteria in liquid culture and monitored the mRFP expression over time at 37°C and 18°C (Figure 18 a and b).

**Figure 18:**
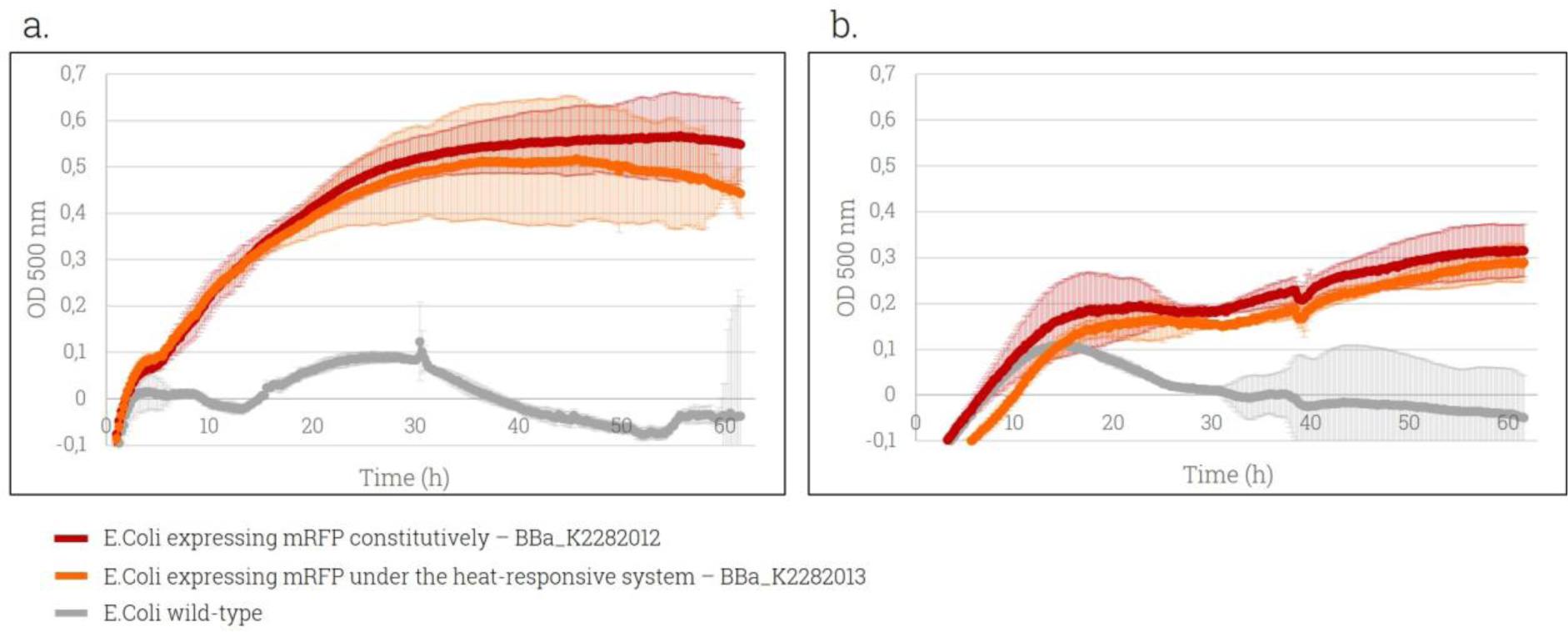
Expression of mRFP induced by the heat-responsive plasmid compared to the positive control over time. **a.** OD500nm given by pSB1C3-BBa_K2282012 (constitutive mRFP expression) and pSB1C3-BBa_K2282013 (heat-induced mRFP expression) at 37°C; **b.** OD500nm given by pSB1C3-BBa_K2282012 (constitutive mRFP expression) and pSB1C3-BBa_K2282013 (heat-induced mRFP expression) at 18°C.

**Fig. 19:**
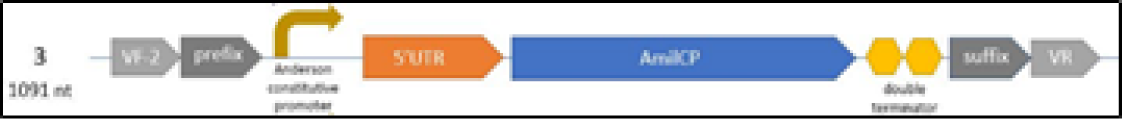
Anderson Constitutive promoter (BBa_J23100) - 5’UTR (BBa_K2282002) - AmilCP (BBa_K592009) - Double terminator (BBa_B0015)

As expected, the graph obtained at 37°C shows that mRFP was expressed at a similar level under the expression of both genetic constructions. The results at 18°C globally show a lower protein expression, explained by a non-optimal growth temperature. Surprisingly, both curved do not differ and suggest a same expression level in presence and absence of the cI857 repressor, in accordance with the previous results obtained on petri dishes. The considerable standard deviations and the apparent lowering of the wild-type DH5-α curve was probably due to little evaporation of the culture medium.

**Fig. 20:**
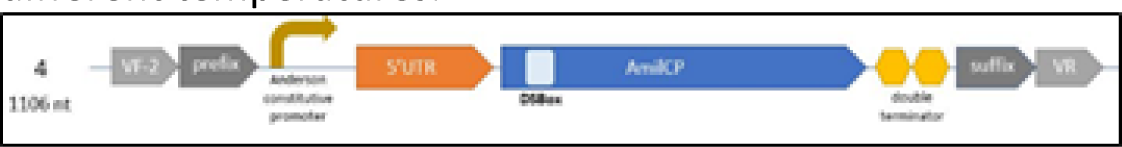
Anderson constitutive promoter (BBa_J23100) - 5’UTR (BBa_K2282002) - AmilCP & DS Box (BBa_K2282001) - Double Terminator (BBa_B0015)

## Discussion

### Cold response

In this article we demonstrated a proof of concept using the blue color reporter amilCP that our cold-inducible system works as expected. Even though the results we have are limited because we relied on a colorimetric assay, this shows the potential of the mechanism for many applications. The temperature starting point of the high expression activation seems to be around 20°C. This is interesting regarding scientific literature since most of the articles we reviewed about Cold-shock proteins (CSPs) relies on the 15°C threshold rather than the 20°C one [REF 3]. The appearance of the gradient indicates clearly that the mechanism is not a strict switch mechanism but a progressive one. The modified mRNA of amilCP shows high stability at low temperature, low stability at high temperature, and an intermediate stability between both. What we wanted to achieve with the CSP mechanism is the deletion of the protein expression under its regulation at a temperature close to the activating temperature of our heat response (30°C). At 27°C, the grey color shows low expression of AmilCP and gives us promises about the validation of our strategy.

**Fig. 21:**
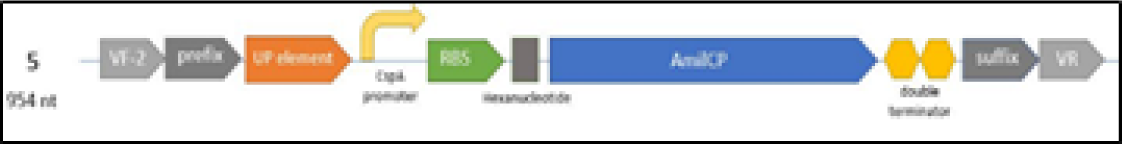
UP element + CspA promoter (BBa_K2282003) - RBS (BBa_B0034) - AmilCP (BBa_K592009) - Double terminator (BBa_B0015)

**Fig. 22:**
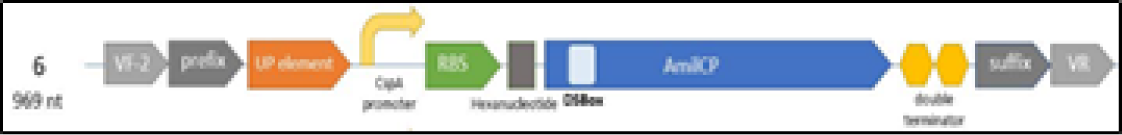
UP element + CspA promoter (BBa_K2282003) - RBS (BBa_B0034) - AmilCP & DS Box (BBa_K2282001) - Double terminator (BBa_B0015)

The promises given by our results to validate the CSPs mimicry as being a reliable mechanism for cold-inducible protein expression are interesting. Perspectives of characterization for the future are numerous. We designed several parts for improving the characterization of our cold-response system. Although we did not have the time to use them in the iGEM competition, they can potentially be used by other teams:

#### BBa_K2282007

to check the influence of 5’-UTR (5’ Untranslated Region) on the amilCP expression.

#### BBa_K2282008

to check if both DS Box and 5’-UTR have an influence on the amilCP expression at different temperatures.

#### BBa_K2282009

to check the influence of the UP element and CspA promoter on the amilCP expression (at a transcriptional level)

#### BBa_K2282010

Checking the influence of the UP element, the CspA promoter and the DS Box on the amilCP expression

These parts can provide a solid background concerning the behavior of each elements of the Cold-shock protein (CSP) system when characterized separately and together with the DS box. What could also be interesting is to use a Green Fluorescent Protein (GFP) rather than amilCP for protein quantification. We chose amilCP in our lab due to the lack of a fluorimeter.

Once the system is completely characterized, the next step would be to replace the reporter gene in the construction by a protein of interest to be expressed at low temperature only. This will require further modeling as for the effect of the Downstream (DS) box insertion in the new protein and its expression at low temperature as well. Based on the protein of interest, a Type III expression system might also be needed to enable secretion of the protein. This will also need characterization because secretion system at low temperature might be difficult to implement for a given protein.

### Heat response

The efficiency of the complete heat-inducible genetic construct created here was also evaluated. The results obtained at 37°C shows that monomeric red fluorescent protein (mRFP) under the expression of the pL promoter was expressed at a similar level with or without its thermo-labile cI857 repressor. This is consistent with what we expected, as the pL promoter is constitutively activated in both cases at this temperature. Indeed, at this temperature, cI857 is denatured and cannot dimerize. It can no longer bind the pL promoter, allowing a regular mRFP expression [REF 17].

The results at 18°C are not consistent with our theoretical expectations, and confirm the visual results previously obtained. Indeed, the heat-shock plasmid induced a similar mRFP expression compared to the positive control, whereas this genetic construction was supposed to trigger gene expression above 37°C only. This lack of cI857 inhibition could have several explanations. First, the BioBrick construction itself could be questioned. The transcription of mRFP may have induced a conformational change in the double terminator we used, preventing the transcription of cI857 or considerably lowering it to a point where the repressor cannot perform its inhibitory action anymore.

As the two ribosome binding sites (RBS) used are the same, we could also question the relative strength of both promoters used. The promoter used for cI857 expression comes from a small combinatorial library of constitutive promoters derived from a consensus sequence by Chris Anderson. Among the different promoters existing in the collection, displaying variable strengths, we chose BBa_J23100 which induces the highest relative expression after the consensus promoter (iGEM Registry of Standard Biological Parts). The challenge is then to compare its relative strength with the pL promoter, derived from a phage, in standardized units. To test this hypothesis, we could design a new sequence coding for the mRFP expression under the Anderson promoter used. Measuring the relative expression of the same reporter gene under both promoters will allow us to assess if a significant difference can be observed.

Finally, this result could come from a lower affinity of the Anderson promoter for the RNA polymerase. Its systematic binding to the pL promoter would explain the absence of difference between our constitutive control and the sequence in which the repressor has been added. Several axes of research are then opened to test our hypothesis: we could first use another promoter for the expression of cI857, known to have a similar affinity for the RNA polymerase as the pL promoter. An alternative would consist in constructing another BioBrick by using standard iGEM parts and test the same construction with the pR promoter. Indeed, the system based on pR promoter regulated by cI repressor under the Anderson BBa_J23114 promoter has already been successfully tested by previous teams (Part BBa_K608351).

The negative results obtained from our heat-inducible system are not a significant drawback for Thermo-responsive plasmid (TRP) development. The system using the pL/pR promoters with the cl857 repressor has already been widely characterized and there are several possible reasons for its failure in our experiments: our genetic construction might have simply been wrong: the reverse orientations of both cl857 and mRFP genes might have accounted for the negative results. What is possible is to test the system with a clean BioBrick standard assembly R10, using cl857 and an mRFP gene regulated by either pL or pR. If the mechanism persists in being non-functional, we could test on other high temperature induced mechanisms. For example, the 5’ Untranslated Transcribed Region (UTR) of *P. aeruginosa* virulence factors has been shown to regulate its mRNA stability, much like the 5’UTR of Cold-shock protein A (CspA) but at high temperature [REF 30].

In summary, one of the two mechanisms tested worked as expected and we provided a great basis for other projects to rely on for the future. Many things need to be done but we are confident that the TRP development can be achieved and used for many different applications.

Each year plants are subjected to temperature stresses such as ice crystal formation at low temperatures and desiccation at high temperature, leading to yield and economic loss. Transforming a microorganism with the TRP to make it express a specific protectant at the plant surface depending on the outside temperatures could be of great value.

Below 15°C, ice-binding proteins will interact with ice crystals to either inhibit their growth (antifreeze proteins) or favoring the nucleation process (ice-nucleation proteins). Despite their opposite functions, both strategies could be of great help to prevent frost damage in their own way and small-scale tests are required to make a final choice. Above 37°C, light-reflecting compounds will limit evapotranspiration by creating a reflective layer. Once applied on crops, the solution will possess a double protection: anti-drought and anti-frost (Figure 23).

**Fig. 23:**
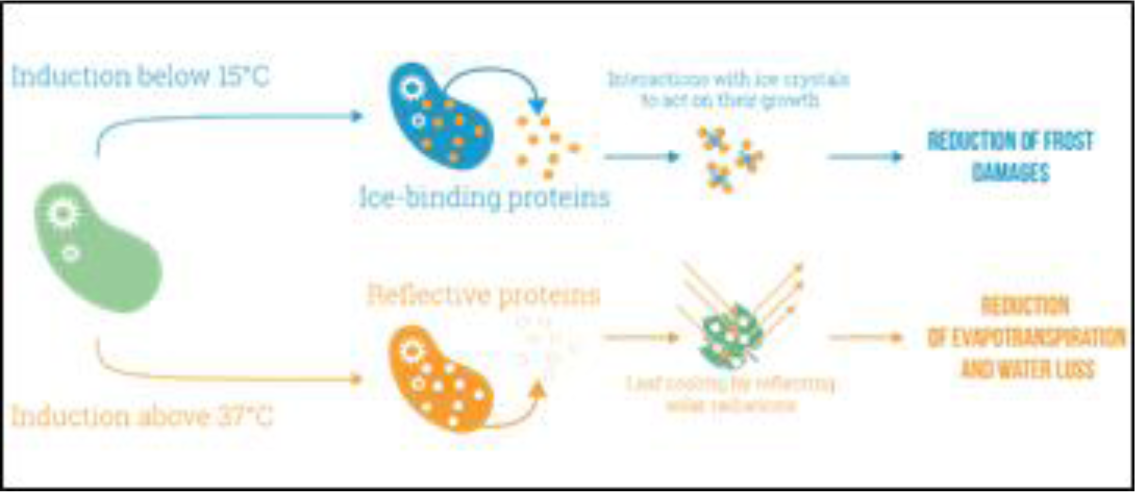
TRP plant application graphic

Plants record the daily average temperature and store it in a total accumulated sum called “thermal time”. As soon as the needed thermal time is reached, the plant can go to the next stage of development. The challenge of recording a total of heat accumulated over time, instead of a temperature absolute value, could be addressed by our bi-functional plasmid. An organism could be transformed with TRP to accumulate a red pigment above a certain temperature in an irreversible manner. A red progression would then be visible as a function of the heat recorded over time. Being able to visualize thermal time could be useful to predict crop development in a particular location or retro-calculate the best sowing date. Alternatively, we could use a TRP-transformed organism to record the average temperature every day, by comparing the color obtained to a reference scale of colors (like pH paper). Directly visualizing the temperature recorded at the plant scale could allow a highly precise evaluation of the potential variations of temperature average within a crop field. Alternatively, we could use our organism in a patch such as the company Cryolog. The company uses organisms in a patch attached to frozen food during its transport. If the cold-chain is broken during the transport of the product, the patch irreversibly turns red. We could inspire ourselves from this system and engineer patches that we could spread across any crop field. If a critical temperature is reached, the TRP will be triggered and a color (blue for low temperature and red for high temperature) will be displayed. It would be a great way to monitor whether the crops have been subjected to critical temperatures. This could furnish an efficient way of monitoring temperature in open fields with contained microorganisms.

Thermoregulation solutions can be costly to implement for industrial or research applications. What could be interesting is to make a thermo-responsive organism induce an exothermic reaction as the temperatures decrease, to reheat the medium, and an endothermic reaction as the temperatures increase in order to cool it down.

## Ethics statement

N/A

## Data availability

All data are fully available without restriction.

