## Supplementary Materials for "A thermo-responsive plasmid for biconditional protein expression"

Response to Reviewers

**Igallagher: Hi IONIS, Thanks for your submission! Please can you add author affiliations.**
→ The engineering school Sup’Biotech was added as the affiliated institution for all authors.

**Igallagher: Some of your references are bold and some are not, this is a small point but keeping consistency in your formatting will ensure it won't distract the reader so we'd suggest modifying these.**
→ The formatting of references was unified across the article.

**Djcamenares: It was a good idea to take into account the effect expression might have on a OD600 reading. Coupled with your OD800 readings you may want to get a count of CFU/mL as a way to verify your conclusions about the amount of bacterial growth.**
→ Calculating the CFU/mL would be an interesting way to confirm the relevance of our OD800 results. However, we did not have access to a proper microscope to count the colonies directly from the culture medium. In order to complement our data, a useful perspective would be to repeat the same experiment and cultivate samples of each conditions to cultivate them on solid agar plates and count the colony quantity.

**I like the presentation of these results. However, either here or in the methods, you should clarify if these pellets represent equal or unequal amounts of bacteria (i.e. did you pellet equal volumes of culture, or equal amounts of bacteria based on OD?)**
→ Indeed, some information missed in the explanation of our methodological approach. We added some details in the methods section as well as in the result description to confirm that equivalent amounts of bacteria were collected for each temperature condition. To do this, we collected equal medium volumes once reaching similar OD600. This explains why we can consider that the differences observed in coloration correspond to differential amilCP expression per bacteria, and exclude the possibility that they are linked to different bacteria quantities.

**Djcamenares: Thoughtful consideration given your lack of equipment (something our team also suffered from). You might be able to supplement your visual observations by actually measuring the color values (RGB) of the pixels in your picture.**
→ Great suggestion ! We extracted the Blue intensity profiles of each pellets on the ImageJ software, and compared their relative levels. Despite aberrant results obtained for one positive control, and imprecision due to poor image quality and pellet inhomogeneity, the results overall appear coherent with the visual observations and can serve to consolidate our results. We added the figure and result analysis, as well as a paragraph about the method used.

**PreethaG: Is there a reason this text is bolded?**
→ No reason, we unified the text format to match the rest of the article.

**The "two plasmid" solution is an interesting one, but it is not clear how this would actually help - you still will have two promoters in your design (and many copies of each in the cell either way if they are on high-copy number plasmids). Perhaps clarify this further in the text or with a diagram**
→ Interesting remark. We thought about this solution as a research tool, to create two “test” plasmids and evaluate the potentiality of our hypothesis towards the results of our heat-shock system. This idea was not to co-transform the two plasmids in the same cells, but testing separately each of them and compare the intensity of the expression obtained. However, after

having discussed it, we came to the conclusion that this method would not necessarily be the most relevant choice.

**Several times the reverse orientation of the two genes/promoters in the TRP has been alluded two (and is probably a good idea) but the construction of this plasmid, with a diagram showing the reverse orientation, is lacking (at least, not presented with the same thoroughness and clarity as the cold-responsive system).**
→ Right, we forgot to include a diagram for the heat-responsive genetic construction as we did for the cold-responsive part. A new one was added at the beginning of the result section for the heat part, showing the complete part we designed, its biobricks and their orientation. This should make our hypothesis towards the impact of the reverse orientation of the genes and promoters easier to visualize.

**Djcamenares: These are good ideas, although reflecting light might not be the best strategy to protect a photosynthetic organism (depending on where the bacteria are applied, of course)**
→ Indeed, several approaches can be considered when dealing with heat protection depending on the targeted effect (water stress, UV damages... ). For safety and regulation problematics, we decided to limit the protective action of our microorganisms to the plant surface only. In this context, it appeared easier to focus on reducing UV damage by reflecting solar radiation rather than acting on internal water stress which would probably require an endogenous action. This strategy turns out to be useful for fruits grown on trees, especially subject to sunburns and premature ripening.

**Djcamenares: A temperature bio-sensor is a great idea, but you might want to reflect on whether or not this application offers any advantages to current methods of measuring and recording temperature (which, I would imagine, are already quick and inexpensive). Perhaps you can highlight that this can show if there is variation of average temperature across a field of crops.**
→ Thank you for this interesting remark. We tried to re-formulate the explanation of our idea, focusing more on the added value compared to existing tools to monitor temperature. In addition to allowing a visual evaluation of temperature variations across a field, the idea of associating a particular color to the amount of heat accumulated in time allows more than the measurement of an absolute value. Even if this will be less precise in term of output, it will allow an irreversible indicator showing that a certain level of heat has been recorded by plants over a certain period regardless of the potential repetitive variations.
